## Supplementary Information for "Precision engineering of biological function with large-scale measurements and machine learning"

Drew S. Tack[1], Peter D. Tonner[1], Abe Pressman[1], Nathanael D. Olson[1], Sasha F. Levy[2,3], Eugenia F. Romantseva[1], Nina Alperovich[1], Olga Vasilyeva[1], David Ross[1]*.

[1]National Institute of Standards and Technology, Gaithersburg, MD, 20899, USA.

[2]SLAC National Accelerator Laboratory, Menlo Park, CA, 94025, USA

[3]Joint Initiative for Metrology in Biology, Stanford, CA, 94305, USA

### Contents

S1. Mutations used for ML-enabled forward engineering .................... 2

S2. *In silico* selection with only single-mutant data .................... 3

### S1. Mutations used for ML-enabled forward engineering

To demonstrate ML-enabled forward engineering, we used a restricted set of 16 missense mutations, chosen to give a range of different effects on the dose-response and to be distributed across the LacI core domain (Fig 7, Table S1).

**Table S1. Mutations used for ML-enabled forward engineering**

| Mutations | Characteristics |
|---|---|
| S69T, S70R, V80L, V136E | Mutations near the ligand pocket, but with different predicted effects |
| I83M, L169P | Mutations with similar effect on the dose-response, but in different regions of the protein structure: I83M is at the dimer interface in the N-terminal core subdomain; L169P is in the periphery of the C-terminal core subdomain. |
| I83F | At same position as I83M, but different effect on dose-response |
| S97T, S279T | Mutations with similar effect on the dose-response, but in different regions of the protein structure: S97T is at the dimer interface in the N-terminal core subdomain; S279T is near the dimer interface in the C-terminal core subdomain. |
| F161V, G200C | Mutations with similar effect on the dose-response, but in different regions of the protein structure: F161V is near ligand pocket and part of the core-pivot region [citation]; G200C is in the protein periphery far from the ligand pocket. |
| Q181H, S190C, N208Y, Q231R, V271M | Mutations that are silent in wild-type background |

### S2. *In silico* selection with only single-mutant data

To assess the importance of multi-mutant data for *in silico* selection, we repeated the multi-objective selections (Figs 3-4) using a subset of the large-scale dataset: LacI variants with only one missense mutation. For each specification, we compared the number of variants with median estimated Hill equation parameters meeting the specification for the full dataset vs. the single-mutant subset. We also compared the maximum probability of success based on the posterior samples reported in the dataset. The results are shown in Table S1. For every specification, the number of variants with estimated Hill parameters meeting the specification and the maximum probability of success were both lower with the single-mutant subset of the data, compared to results with the full dataset. Also, many of the targeted specifications were not accessible by any variants in the single-mutant dataset. The detailed quantitative criteria for each specification and the method of calculating the probability of success are described in the Materials and Methods.

**Table S2. Comparison of in silico selection with full dataset vs. single-mutant dataset.**

| | Full dataset | | Single-mutant dataset | |
|---|---|---|---|---|
| Specification | Number of variants | Max success probability | Number of variants | Max success probability |
| $EC_{50}$ = 10 µmol/L<br>$G_{\infty}$ = 16 kMEF | 18 | 0.41 | 0 | N/A |
| $EC_{50}$ = 30 µmol/L<br>$G_{\infty}$ = 16 kMEF | 61 | 0.63 | 0 | N/A |
| $EC_{50}$ = 100 µmol/L<br>$G_{\infty}$ = 16 kMEF | 109 | 0.66 | 2 | 0.31 |
| $EC_{50}$ = 10 µmol/L<br>$G_{\infty}$ = 25 kMEF | 407 | 0.66 | 15 | 0.47 |
| $EC_{50}$ = 30 µmol/L<br>$G_{\infty}$ = 25 kMEF | 1513 | 0.75 | 45 | 0.66 |
| $EC_{50}$ = 10 µmol/L<br>Inverted dose-response | 1 | 0.47 | 0 | N/A |
| $EC_{50}$ = 30 µmol/L<br>Inverted dose-response | 1 | 0.34 | 0 | N/A |
| $EC_{50}$ = 100 µmol/L<br>Inverted dose-response | 2 | 0.19 | 0 | N/A |
